# Ensemble Feature Selection and Meta-Analysis of Cancer miRNA Biomarkers

**DOI:** 10.1101/353201

**Authors:** Lopez-Rincon Alejandro, Martinez-Archundia Marlet, Martinez-Ruiz Gustavo Ulises, Tonda Alberto

**Affiliations:** Life Sciences and Health, CWI, Amsterdam, Netherlands; Laboratorio de Modelado Molecular, Bioinformática y Diseño de Fármacos. Escuela Superior de Medicina, Instituto Politecnico Nacional, Mexico City, Mexico; Federico Gomez Children’s Hospital; School of Medicine, National Autonomous University of Mexico, Mexico City, Mexico; UMR 782 GMPA, Université Paris-Saclay, INRA, AgroParisTech, Paris, France

## Abstract

The role of microRNAs (miRNAs) in cellular processes captured the attention of many researchers, since their dysregulation is shown to affect the cancer disease landscape by sustaining proliferative signaling, evading program cell death, and inhibiting growth suppressors. Thus, miRNAs have been considered important diagnostic and prognostic biomarkers for several types of tumors. Machine learning algorithms have proven to be able to exploit the information contained in thousands of miRNAs to accurately predict and classify cancer types. Nevertheless, extracting the most relevant miRNA expressions is fundamental to allow human experts to validate and make sense of the results obtained by automatic algorithms. We propose a novel feature selection approach, able to identify the most important miRNAs for tumor classification, based on consensus on feature relevance from high-accuracy classifiers of different typologies. The proposed methodology is tested on a real-world dataset featuring 8,129 patients, 29 different types of tumors, and 1,046 miRNAs per patient, taken from The Cancer Genome Atlas (TCGA) database. A new miRNA signature is suggested, containing the 100 most important oncogenic miRNAs identified by the presented approach. Such a signature is proved to be sufficient to identify all 29 types of cancer considered in the study, with results nearly identical to those obtained using all 1,046 features in the original dataset. Subsequently, a meta-analysis of the medical literature is performed to find references to the most important biomarkers extracted by the methodology. Besides known oncomarkers, 15 new miRNAs previously not ranked as important biomarkers for diagnosis and prognosis in cancer pathologies are uncovered. Such miRNAs, considered relevant by the machine learning algorithms, but still relatively unexplored by specialized literature, could provide further insights in the biology of cancer.

## Author summary

MicroRNAs (miRNAs) are non-coding RNA molecules that regulate gene expression. In the last years, the under and over expression of miRNAs has been related to the diagnosis and prognosis of specific cancer types. While machine learning techniques can efficiently exploit the information contained in thousands of miRNAs to detect the presence and typology of tumors, it is still fundamental to isolate the minimum possible number of meaningful features, in order to allow human experts to validate the results. We propose a new ensemble feature selection methodology, and we test it on a real-world dataset, taken from The Cancer Genome Atlas (TCGA) database. The considered dataset contains 1,046 miRNA expressions, data for 8,129 patients, with 29 classes of tumors. Feature selection is performed by considering the 100 most relevant features emerging from the consensus between 8 state-of-the-art classifiers with high accuracy on the dataset. Such list is shown to be sufficient to provide an unchanged classification accuracy. Finally, the 50 most important features selected by our approach are validated by human experts, resorting to a literature review. Interestingly, while most of the selected miRNAs are known oncomarkers, a few appear still understudied, and might thus represent promising leads for future research.

## 1 Introduction

Several studies have shown the properties of microRNA types (miRNAs) as oncogenes and tumor suppressors [1–3]. Since then, many sophisticated techniques, such as high-throughput technologies, microarray, mass spectrometry and especially the Next Generation Sequencing (NGS), have been developed for their identification [4]. However, it is clear that the development of computational tools is needed for the interpretation of results from these high-throughput experiments [5]. Indeed, computational assisted methods are used for the identification of miRNAs from different genome organisms, for example in *Caenorhabditis briggsae* [6] and in Epstein-Barr virus (EBV or HHV4), a member of the human herpesvirus (HHV) [7]. Furthermore, several computational techniques can be applied to accurately predict miRNA expressions, as seen for example in [8].

Succeeding the earliest evidence of miRNA involvement in human cancer by Croce and collaborators [9], various studies demonstrate that miRNA expression is deregulated in human cancer through diverse mechanisms [10]. Additionally, in comparison to the impractical and invasive methods currently used for cancer diagnosis [11,12], miRNA biomarkers can be detected directly from biological fluids (such as blood, urine, saliva and pleural fluid [13]), and they can also be used as biomarkers to detect tumors at an early stage, which is extremely important for survival. For example, the 5-year survival rate for lung cancer is 5%, but an early diagnosis can boost it to almost 50% [14]. Thus, miRNA expression profiles correlate with clinical variables, highlighting their potential value as prognostic and/or diagnostic tools.

In such a context of increasing availability of data, it is of utmost practical importance to build databases of miRNA expressions data for cancer research [15–19], and also to extract features that could be used as cancer biomarkers [20–22]. For example, miRNA *hsa-mir-21* is mentioned as a marker for patients with squamous cell lung carcinoma [23], with astrocytoma [24], breast cancer [25], and gastric cancer [26]. Following this idea, the scientific community is currently looking for miRNA signatures, representing the minimal number of miRNAs to be measured for discriminating between different stages and types of cancer.

Current NGS technologies such as Applied Biosystems, SOLiD3,or HiSeq from Illumina are able to extract thousands of components in genome sequences [27], and traditional linear statistical analysis are not suited to manage such quantities of measured elements with non-lineal relationships to extract meaningful features. Thus, a suitable solution, is to use machine learning techniques for analysis, classification, and relevant features extraction of miRNA data [28–30].

Starting from a dataset containing 8,129 patients, 29 different types of cancer, and 1,046 different miRNA expressions, 8 state-of-the-art classifiers are used to extract the most relevant miRNAs to use as biomarkers for cancer classification. Typically, classifiers trained on a dataset will not use the whole set of available features to separate classes, but just a subset which could be ordered by relative importance, with a different meaning given to the list by the specific technique. The top 100 biomarkers in the list are then evaluated as a potential reduced signature for classification. Finally, the top 50 miRNAs are compared to a meta-analysis of the medical literature, to validate the results automatically produced by the machine learning algorithms. Unsurprisingly, most of the miRNAs identified by the classifiers are also considered important by the specialized literature: 15 of them, however, are still understudied, and they could thus represent promising leads for future exploration.

The rest of the paper is organized as follows. The target dataset and the proposed approach are detailed in Section 2. Experimental results are reported in Section 3, while Section 4 concludes the paper.

## 2 Methods

The considered dataset, containing miRNA sequencing isoform values, is taken from the Cancer Genome Atlas^1^. The database contains the information from 8,129 patients. Using the next-generation sequencing miRNASeq BCGSC IlluminaHiSeq miRNASeq Level_3, a total of 1,046 miRNA expression features for each case study are extracted. In summary, the dataset that will be used in the following experiments has 29 types of tumors, 1,046 miRNA features, and 8,129 patient samples. Information on the dataset is summarized in Table 1.

**Table 1.** Dataset: Tumor type, class label, and number of samples per class.

| Tumor Type | Number of Samples | Class |
| --- | --- | --- |
| Adrenocortical carcinoma [ACC] | 80 | 0 |
| Bladder Urothelial Carcinoma [BLCA] | 415 | 1 |
| Breast invasive carcinoma [BRCA] | 778 | 2 |
| Cervical squamous cell carcinoma [CESC] | 308 | 3 |
| Cholangiocarcinoma [CHOL] | 35 | 4 |
| Esophageal carcinoma [ESCA] | 47 | 5 |
| FFPE Pilot Phase II [FPPP] | 200 | 6 |
| Head and Neck squamous cell carcinoma [HNSC] | 45 | 7 |
| Kidney Chromophobe [KICH] | 488 | 8 |
| Kidney renal clear cell carcinoma [KIRC] | 66 | 9 |
| Kidney renal papillary cell carcinoma [KIRP] | 261 | 10 |
| Lower Grade Glioma [LGG] | 292 | 11 |
| Liver hepatocellular carcinoma [LIHC] | 530 | 12 |
| Lung adenocarcinoma [LUAD] | 374 | 13 |
| Lung squamous cell carcinoma [LUSC] | 458 | 14 |
| Lymphoid Neoplasm Diffuse Large B-cell Lymphoma [DLBC] | 341 | 15 |
| Mesothelioma [MESO] | 87 | 16 |
| Pancreatic adenocarcinoma [PAAD] | 179 | 17 |
| Pheochromocytoma and Paraganglioma [PCPG] | 184 | 18 |
| Prostate adenocarcinoma [PRAD] | 500 | 19 |
| Sarcoma [SARC] | 262 | 20 |
| Skin Cutaneous Melanoma [SKCM] | 452 | 21 |
| Stomach adenocarcinoma [STAD] | 399 | 22 |
| Testicular Germ Cell Tumors [TGCT] | 155 | 23 |
| Thymoma [THYM] | 514 | 24 |
| Thyroid carcinoma [THCA] | 124 | 25 |
| Uterine Carcinosarcoma [UCS] | 418 | 26 |
| Uterine Corpus Endometrial Carcinoma [UCEC] | 57 | 27 |
| Uveal Melanoma [UVM] | 80 | 28 |

As a baseline comparison, a preliminary analysis of the available data is performed, normalizing all the isoform expressions altogether and then quantifying the highest expressed miRNAs for each cancer tumor type. Next, the top 50 most expressed miRNAs for each tumor type are arranged in descending order. Finally, a coefficient 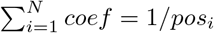 where *N* is the number of tumor types in the dataset, that depends on miRNA’s relative position, bringing the result displayed in Figure 1.

**Fig 1.**
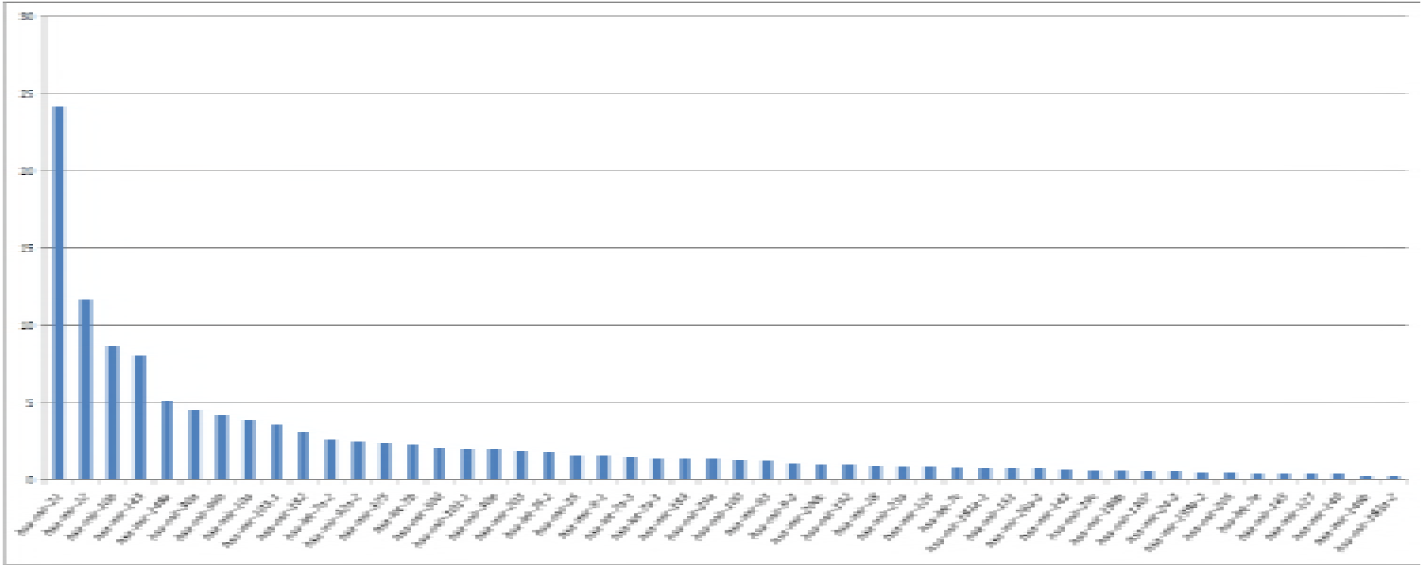
Top 50 most expressed miRNA types, across all cancer classes in the considered dataset.

As the objective is to find and validate a reduced list of miRNAs to be used as a signature, feature selection is to be performed on the dataset. Popular approaches to feature selection range from univariate statistical considerations, to iterated runs of the same classifier with a progressively reduced number of features, in order to assess the contribution of the features to the overall result. As the considered case study is particularly complex, however, relying upon simple statistical analyses or a single classifier might not suffice. Following the idea behind *ensemble feature selection* [31–33], we use multiple algorithms to obtain a more robust predictive performance. For this purpose, we train a set of classifiers to then extract a sorted list of the most relevant features from each. As, intuitively, a feature considered important by the majority of classifiers in the set is likely to be relevant for our aim, the information from all classifiers is then compiled to find the most common relevant features.

Starting from a thorough comparison of 22 different state-of-the-art classifiers on the considered dataset presented in [34], in this work a subset of those classifiers is selected considering both (i) high accuracy and (ii) a way to extract the relative importance of the features from the trained classifier. After preliminary tests to set algorithms’ hyperparameters, 8 classifiers are chosen, all featuring an average accuracy higher than 90% on a 10-fold cross-validation:

- **`BaggingClassifier`** [35]
- **`GradientBoostingClassifier`**[36]
- **`LogisticRegression`** [37]
- **`PassiveAggressiveClassifier`** [38]
- **`RandomForestClassifier`** [39]
- **`RidgeClassifier`** [40]
- **`SGDClassifier`** (Stochastic Gradient Descent on linear models) [41]
- **`SVC`** (Support Vector Machines Classifier with a linear kernel) [42]

All considered classifiers are implemented in the **`scikit-learn`** Python toolbox [43].

Overall, the selected classifiers fall into two broad typologies: those exploiting ensembles of classification trees [44] (**`Bagging`**, **`GradientBoosting`**, **`RandomForest`**), and those optimizing the coefficients of linear models to separate classes (**`LogisticRegression`**, **`PassiveAggressive`**, **`Ridge`**, **`SGD`**, **`SVC`**). Depending on classifier typology, there are two different ways of extracting relative feature importance. For classifiers based on classification trees, the features used in the splits are counted and sorted by frequency, from the most to the least common. For classifiers based on linear models, the values of the coefficients associated to each feature can be used as a proxy of their relative importance, sorting coefficients from the largest to the smallest in absolute value. As the two feature extraction methods return heterogeneous numeric values, only the relative sorting of features provided by each classifier is considered. We arbitrarily decide to extract the top 100 most relevant features, so we assign to each feature *f* a simple score *S*_*f*_ = *N*_*t*_/*N*_*c*_, where *N*_*t*_ is the number of times that specific features appears among the top 100 of a specific classifier instance, while *N*_*c*_ is the total number of classifiers instances used; for instance, a feature appearing among the 100 most relevant in 73% of the classifiers used would obtain a score *S*_*f*_ = 0.73. In order to increase the generalizability of our results, each selected classifier is run 10 times, using a 10-fold stratified cross-validation, so that each fold preserves the percentage of samples of each class of the original dataset. Thus, *N*_*c*_ = 80 (8 types of classifiers, run 10 times each). The complete procedure is summarized by Algorithm 1.

Finally, the top 50 features obtained in this way are validated with a meta-analysis of the relevant literature. In a first step, reviews of miRNA types related to cancer are inspected for the presence of the extracted features. Subsequently, the PubMed database is interrogated for references containing the identified miRNA types^2^, and the results are later manually analyzed by the authors.

## 3 Results and Discussion

Table 2 compares the classification accuracy of each classifier using the full 1,046 features, with the accuracy obtained by the same classifier using a signature composed by selected 100 features. It is interesting to notice how the accuracy is, for most cases, unchanged, providing empirical evidence that a 100-miRNA signature is enough to obtain good classification results.

**Table 2.**
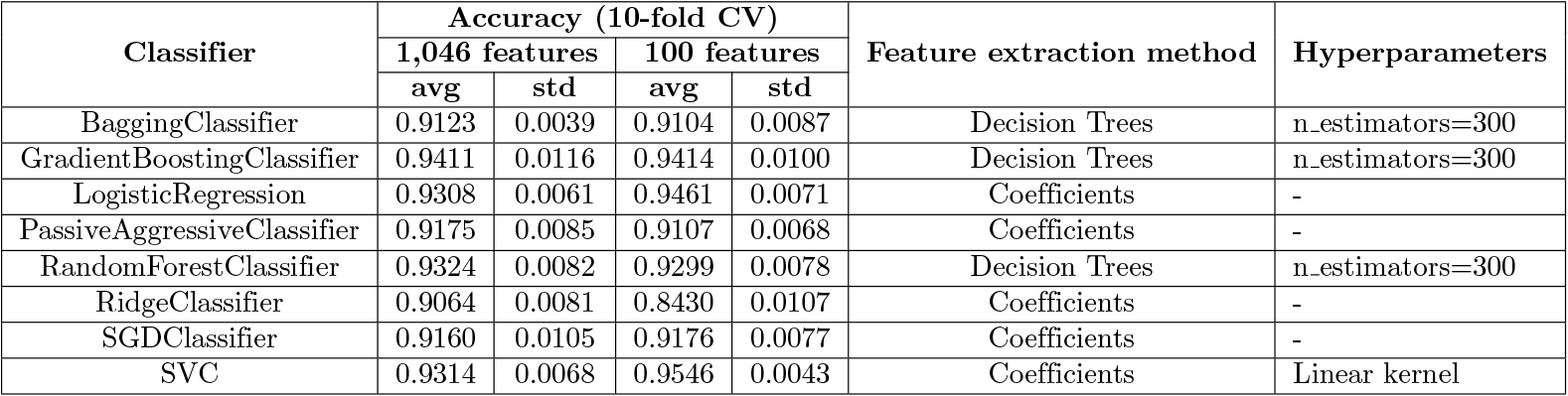
Classifiers used in the experiments. For each classifier, the average accuracy and corresponding standard deviation on a 10-fold cross validation are reported, for both the complete dataset (1,046 features) and the 100 features that have been selected as the most relevant. In the case a classifier is not using standard values for its hyperparameters, the relevant variations are summarized in the corresponding column.

| Classifier | Accuracy (10-fold CV) |  |  |  | Feature extraction method | Hyperparameters |
| --- | --- | --- | --- | --- | --- | --- |
|  | 1,046 features |  | 100 features |  |  |  |
|  | avg | std | avg | std |  |  |
| BaggingClassifier | 0.9123 | 0.0039 | 0.9104 | 0.0087 | Decision Trees | n_estimators=300 |
| GradientBoostingClassifier | 0.9411 | 0.0116 | 0.9414 | 0.0100 | Decision Trees | n_estimators=300 |
| LogisticRegression | 0.9308 | 0.0061 | 0.9461 | 0.0071 | Coefficients | - |
| PassiveAggressiveClassifier | 0.9175 | 0.0085 | 0.9107 | 0.0068 | Coefficients | - |
| RandomForestClassifier | 0.9324 | 0.0082 | 0.9299 | 0.0078 | Decision Trees | n_estimators=300 |
| RidgeClassifier | 0.9064 | 0.0081 | 0.8430 | 0.0107 | Coefficients | - |
| SGDClassifier | 0.9160 | 0.0105 | 0.9176 | 0.0077 | Coefficients | - |
| SVC | 0.9314 | 0.0068 | 0.9546 | 0.0043 | Coefficients | Linear kernel |

Figure 2 shows a heatmap comparing the relative frequency of the overall top 100 most frequent features, for each considered classifier. As expected, not all classifiers use the same features to separate the types of cancer, and thus using their consensus proves to be more robust than just relying upon a single algorithm. It is interesting to notice that while the overall most common biomarkers appear among the top for each classifier, some classifiers make use of only a few. For example, **`BaggingClassifier`** and **`RidgeClassifier`** do not use the vast majority of the features exploited by others to discriminate between classes. A further difference between the two is that features used by **`BaggingClassifier`** that are also appearing in the top 100 are clearly important for the classifier, being used in almost 100% of its 10 runs; while it is noticeable how **`RidgeClassifier`** probably bases its discrimination on features that do not appear among the top 100. This also explains the drop in performance when **`RidgeClassifier`** is forced to use the top 100 features; while **`BaggingClassifier`** seems to be overall unaffected by the restriction (see Table 2). One classifier, **`SVC`**, even slightly increases its average accuracy, probably due to the fact that the search space defined by the 100-feature signature is easier to explore for its optimization procedure.

**uFig 1.**
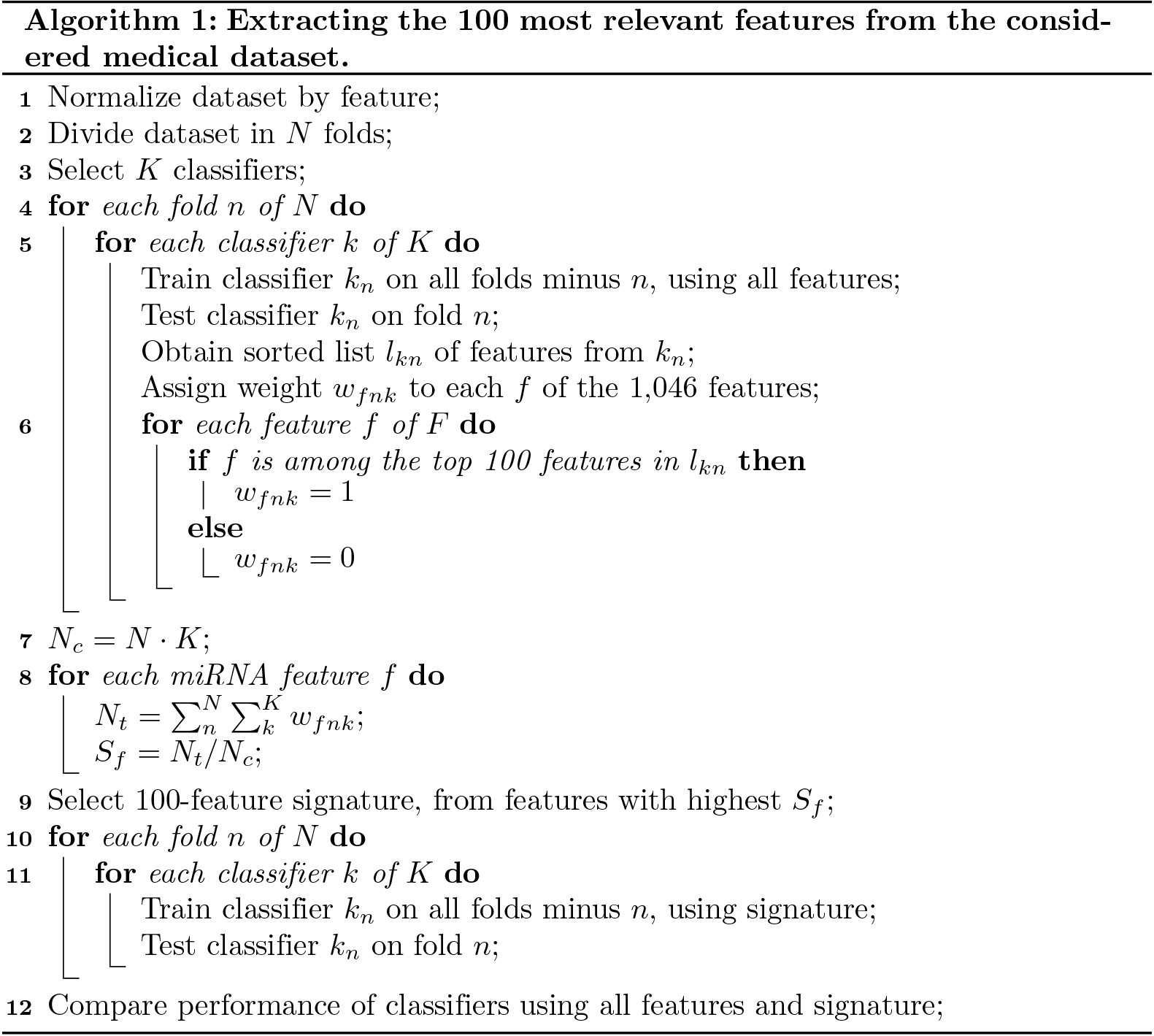

**Fig 2.**
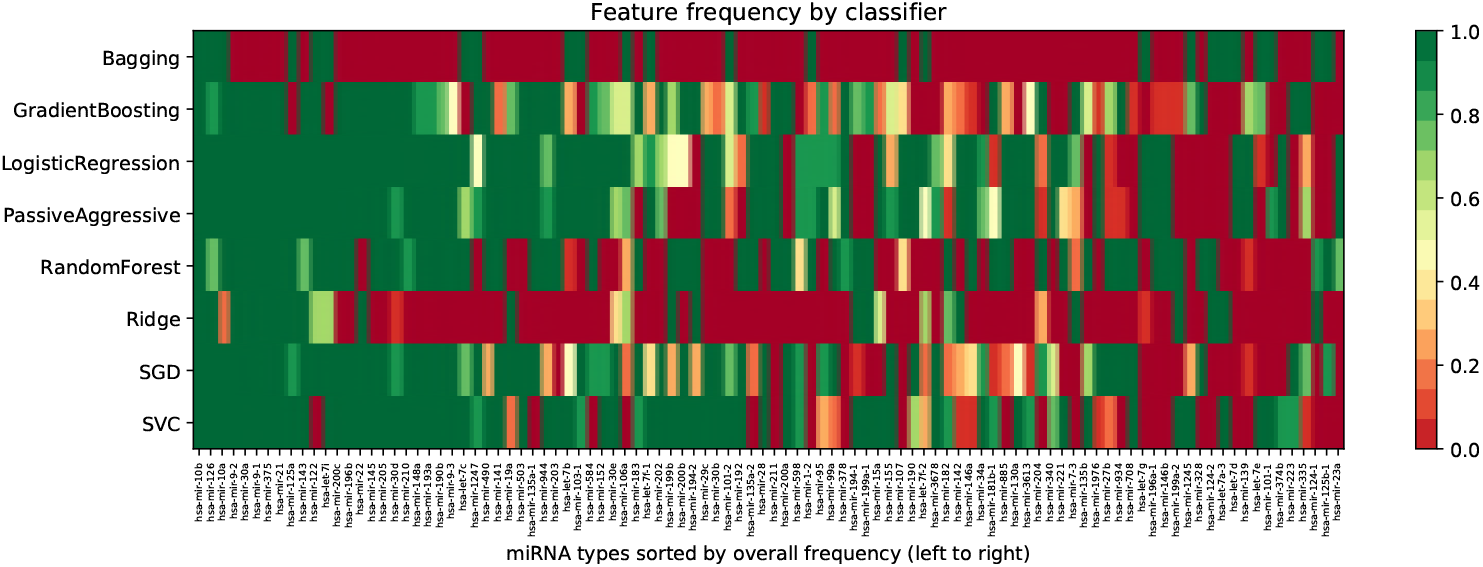
Heatmap with the frequency of the overall top 100 most frequent features, divided by classifier. Features are sorted by frequency, from left to right.

The top 50 features extracted, reported in Table 4, are then validated through a meta-analysis of the specialized literature. The proposed meta-analysis is mainly based on two surveys of miRNA biomarkers, Muller et al. [1], and Ferracin et al. [2], plus 150 papers obtained through a PubMed query and later manually inspected by the authors.

**Table 3.** miRNA types identified by the machine learning feature extraction, appearing in less than 50 references connected to cancer in a PubMed query.

| miRNA type | References related to cancer | References NOT related to cancer |
| --- | --- | --- |
| hsa-mir-9-1 | 36 | 8 |
| hsa-mir-190b | 8 | 9 |
| hsa-mir-9-3 | 20 | 9 |
| hsa-mir-1247 | 14 | 8 |
| hsa-mir-490 | 39 | 20 |
| hsa-mir-135a-1 | 1 | 2 |
| hsa-mir-944 | 16 | 3 |
| hsa-mir-103-1 | 2 | 2 |
| hsa-mir-584 | 23 | 16 |
| hsa-mir-202 | 43 | 51 |
| hsa-mir-199b | 45 | 63 |
| hsa-mir-194-2 | 2 | 3 |
| hsa-mir-101-2 | 8 | 4 |
| hsa-mir-135a-2 | 1 | 1 |
| hsa-mir-28 | 47 | 65 |

**Table 4.**
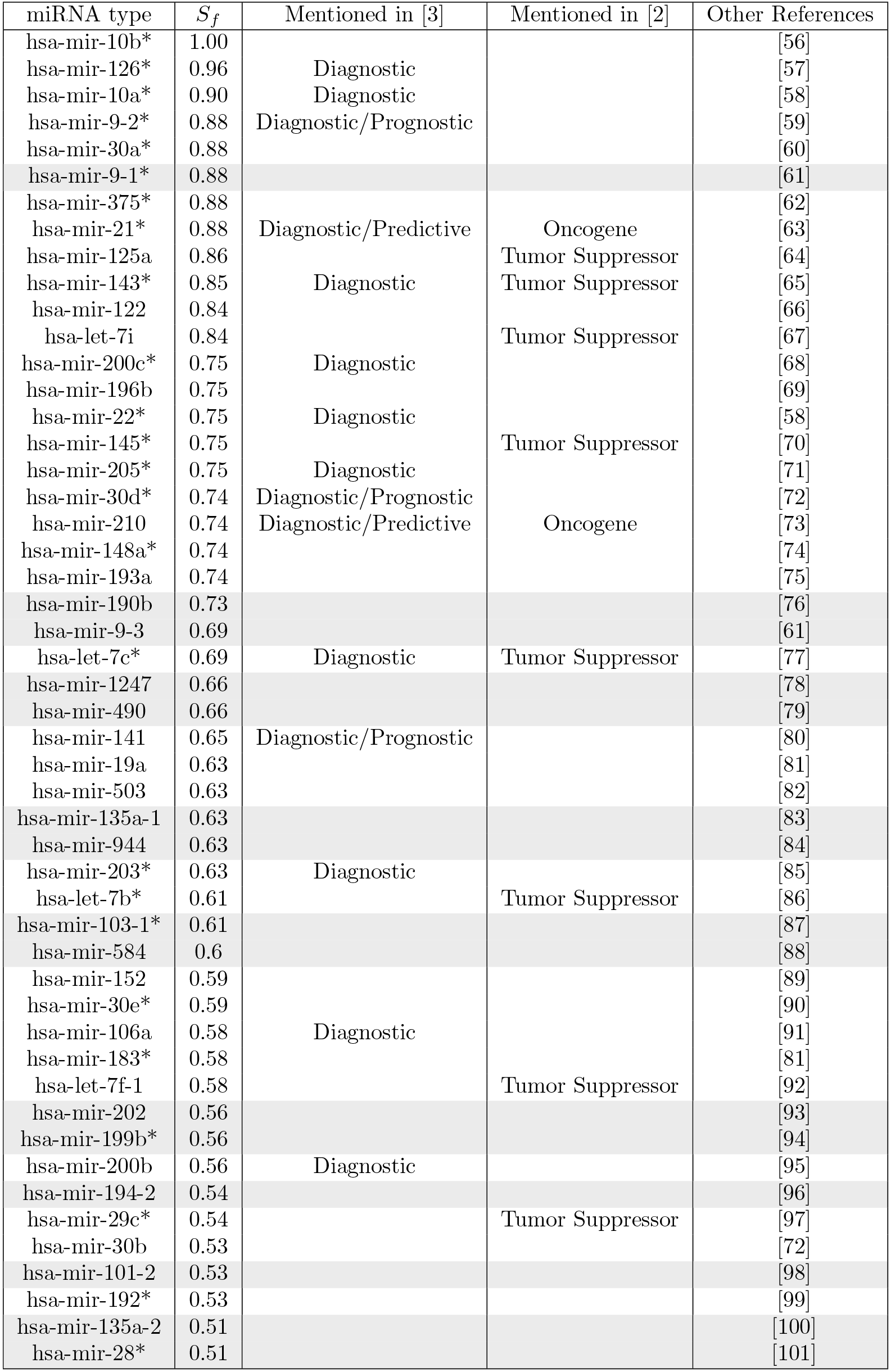
Table comparing the top 50 most frequent features extracted by the machine learning algorithms with existing biomarkers references in literatures. miRNAs highlighted in gray are proposed as promising venues of research, as they only appear in less than 50 references connected to cancer in literature, according to the PubMed query performed in this study. miRNAs marked with * show considerable overexpression along all classes of tumors, as reported in Figure 1.

Of the top 50 features, 15 (*hsa-mir-10a*, *hsa-mir-21*, *hsa-mir-22*, *hsa-mir-30d*, emphhsa-mir-106a, *hsa-mir-126*, *hsa-mir-141*, *hsa-mir-143*, *hsa-mir-200b*, *hsa-mir-200c*, *hsa-mir-203*, *hsa-mir-205*, *hsa-mir-210*, *hsa-mir-375*, and *hsa-let-tc*) are mentioned either for cancer diagnosis, prediction, or prognosis by Muller [3], and can be measured by body fluids. Ferracin [2] mentions a total of 10, with 6 not appearing in [3]: *hsa-let-7b*, *hsa-let-7f-1*, *hsa-let-7i*, *hsa-mir-29c*, and *hsa-mir-145.* In total, 21 miRNAs identified by the proposed approach are found among the two surveys.

The most important result of this work is that several miRNAs types highly ranked by the classifiers appear to be still understudied in literature. Table 3 reports a summary of the literature meta-analysis, from which it clearly emerges that 15 of the miRNA types appear in less than 50 references connected to cancer from the PubMed query: strikingly, *hsa-135-a-1* appears only once, and *hsa-103-1* only twice, while both are considered important by more than 60% of the classifiers. These two miRNAs could thus represent promising leads for future research. Interestingly, only 25 of the miRNA types found by the proposed machine learning approach also appear among the most expressed in the baseline analysis summarized in Figure 1, and only 4 (*hsa-mir-9-1*, *hsa-mir-28*, *hsa-mir-103-1*, and *hsa-mir-199b*) are both overexpressed and understudied, suggesting that simple statistical approaches might not be enough to extract meaningful information from such complex data.

*hsa-mir-21*, mentioned in both surveys, is also the most commonly overexpressed miRNA for all classes of tumors in the statistical analysis summarized by Figure 1. *hsa-mir-10b*, the feature with the highest *S*_*f*_ score, while not appearing in the surveys, is currently under clinical trials as oncomarker for Glioma [45], and it is mentioned in a recent article as possible oncomarker [46]. *hsa-mir-135a-1* and *hsa-mir-135a-2,* located inside chromosomes 3 and 12, respectively, generate the same mature active sequence [47]. *hsa-mir-9-1*, *hsa-mir9-2*, and *hsa-mir9-3*, generate the oncogenic *hsa-mir-9* [48]. *hsa-mir-944* at the moment is known to decrease malignant features in gastric [49], colorectal [50] and endometrial [51] cancers.

We envision that understudied miRNAs could play a key role in the biology of cancer. On the other hand, the presence of single-nucleotide polymorphism (SNP) could affect miRNA biology due to alterations in the miRNA maturation process, their structure, and expression levels. From these, creation and loss of miRNA targeted sites by SNPs is the most inspected area [52,53]. However, the miRNA-mediated oncogenic transcriptional landscape could be a consequence of the SNPs presence in the seed region of mature miRNAs [54], that is involved in the molecular recognition with its targeted mRNAs. Although the presence of SNPs in seed regions seems to be negatively selected [55], in a pathological condition as cancer, it would be interesting to perform further measurements to determine whether any miRNA-related SNPs is contained in miRNA signature we propose. The complete list of the 100 extracted features is in Annex A.

## 4 Conclusions

miRNAs regulate the transcriptional landscape in a fine-tuning way. Alterations in miRNA expression profiles have serious consequences for several diseases, such as cancer. Since ectopic modulation of specific miRNAs could compromise the hallmarks of cancer, it has been proposed that cancerous miRNAs could be modulated by microRNA-based therapies. In this sense, several efforts had been achieved to generate scaffold-mediated miRNA-based delivery systems, thus exploiting the miRNA-mediated therapeutic potential. On the other hand, the altered miRNA expression profile present in cancer could be used as prognostic and/or diagnostic marker. In this respect, it has been stated that several miRNA signatures correlate with clinical outcomes, highlighting the need of a miRNA-based clinical decision-making tool. In this paper, we develop a new machine learning approach to obtain a robust, reduced miRNA signature, from a dataset containing 29 different types of cancer. A further meta-analysis of literature to validate the miRNA signature shows both well-known oncogenic and underestimated miRNA types. The results of this work could potentially be used to uncover new, promising leads of research for a better understanding of miRNA behavior.

Furthermore, personal-directed anti-tumoral therapy could be achieved by measurement of a specific, minimal miRNA signature, as proposed in this work.

## Acknowledgements

The results published here are based upon data generated by The Cancer Genome Atlas Research Network, and the authors would like to thank all specimen donors and research groups that contributed to this project.

## Funding

This work was partially funded by INRA, France; and CONACYT, Mexico.

## Annex A

List of the top 100 most relevant features identified by the proposed methodology, in order of importance: hsa-mir-10b, hsa-mir-126, hsa-mir-10a, hsa-mir-9-2, hsa-mir-30a, hsa-mir-9-1, hsa-mir-375, hsa-mir-21, hsa-mir-125a, hsa-mir-143, hsa-mir-122, hsa-let-7i, hsa-mir-200c, hsa-mir-196b, hsa-mir-22, hsa-mir-145, hsa-mir-205, hsa-mir-30d, hsa-mir-210, hsa-mir-148a, hsa-mir-193a, hsa-mir-190b, hsa-mir-9-3, hsa-let-7c, hsa-mir-1247, hsa-mir-490, hsa-mir-141, hsa-mir-19a, hsa-mir-503, hsa-mir-135a-1, hsa-mir-944, hsa-mir-203, hsa-let-7b, hsa-mir-103-1, hsa-mir-584, hsa-mir-152, hsa-mir-30e, hsa-mir-106a, hsa-mir-183, hsa-let-7f-1, hsa-mir-202, hsa-mir-199b, hsa-mir-200b, hsa-mir-194-2, hsa-mir-29c, hsa-mir-30b, hsa-mir-101-2, hsa-mir-192, hsa-mir-135a-2, hsa-mir-28, hsa-mir-211, hsa-mir-200a, hsa-mir-598, hsa-mir-1-2, hsa-mir-95, hsa-mir-99a, hsa-mir-378, hsa-mir-194-1, hsa-mir-199a-1, hsa-mir-15a, hsa-mir-155, hsa-mir-107, hsa-mir-190, hsa-let-7f-2, hsa-mir-3678, hsa-mir-182, hsa-mir-142, hsa-mir-146a, hsa-mir-34a, hsa-mir-181b-1, hsa-mir-885, hsa-mir-130a, hsa-mir-3613, hsa-mir-204, hsa-mir-340, hsa-mir-221, hsa-mir-7-3, hsa-mir-135b, hsa-mir-1976, hsa-mir-27b, hsa-mir-934, hsa-mir-708, hsa-let-7g, hsa-mir-196a-1, hsa-mir-146b, hsa-mir-199a-2, hsa-mir-1245, hsa-mir-328, hsa-mir-124-2, hsa-let-7a-3, hsa-let-7d, hsa-mir-139, hsa-let-7e, hsa-mir-101-1, hsa-mir-374b, hsa-mir-223, hsa-mir-335, hsa-mir-124-1, hsa-mir-125b-1, hsa-mir-23a.

## Footnotes

1 http://cancergenome.nih.gov/

2 Query performed on January 20th, 2018, on https://www.ncbi.nlm.nih.gov/pubmed/. The query used is **`(<mir-number>[TEXT WORD]) AND ((cancer[TEXT WORD]) OR (tumor[TEXT WORD]))`**.

